# PET-BIDS, an extension to the brain imaging data structure for positron emission tomography

**DOI:** 10.1101/2021.06.16.448390

**Authors:** Martin Norgaard, Granville J. Matheson, Hanne D. Hansen, Adam Thomas, Graham Searle, Gaia Rizzo, Mattia Veronese, Alessio Giacomel, Maqsood Yaqub, Matteo Tonietto, Thomas Funck, Ashley Gillman, Hugo Boniface, Alexandre Routier, Jelle R. Dalenberg, Tobey Betthauser, Franklin Feingold, Christopher J. Markiewicz, Krzysztof J. Gorgolewski, Ross W. Blair, Stefan Appelhoff, Remi Gau, Taylor Salo, Guiomar Niso, Cyril Pernet, Christophe Phillips, Robert Oostenveld, Jean-Dominique Gallezot, Richard E. Carson, Gitte M. Knudsen, Robert B. Innis, Melanie Ganz

**Affiliations:** Neurobiology Research Unit, Rigshospitalet, and Institute of Clinical Medicine, Univ. Copenhagen, Denmark; Department of Psychology, Stanford University, California, USA; Department of Psychiatry, Columbia University, New York, NY 10032 USA; Centre for Psychiatry Research, Department of Clinical Neuroscience, Karolinska Institutet and Stockholm Health Care Services, Stockholm, Sweden; Athinoula A. Martinos Center for Biomedical Imaging, MGH/HST, Charlestown, MA, USA; Intramural Research Program, NIMH, Bethesda, USA; Invicro and Division of Brain Sciences, Imperial College London, London, UK; Centre for Neuroimaging Sciences, King’s College London, London, UK; Department of Information Engineering, University of Padua, Padua, Italy; Amsterdam UMC, location VUmc, department of radiology and nuclear medicine, Amsterdam, NL; Université Paris-Saclay, CEA, CNRS, Inserm, BioMaps, Service Hospitalier Frédéric Joliot, Orsay, France; INM-1, Jülich Forschungszentrum, Germany; Aust. e-Health Research Centre, Commonwealth Scientific and Industrial Research Organisation, Townsville, AU; Centre d’Acquisition et de Traitement des Images, CEA, Paris, France; Inria, Aramis project-team, Sorbonne Université, Institut du Cerveau - Paris Brain Institute - ICM, Inserm, CNRS, AP-HP, Hôpital de la Pitié Salpêtrière, Paris, France; Department of Neurology, University of Groningen, University Medical Center Groningen, Groningen, NL; Wisconsin Alzheimer’s Disease Research Center, Division of Geriatrics, Department of Medicine, University of Wisconsin-Madison School of Medicine and Public Health, Madison, WI, USA; Center for Adaptive Rationality, Max Planck Institute for Human Development, Berlin, Germany; Institute of psychology, Université catholique de Louvain, Louvain la Neuve Belgium; Department of Psychology, Florida International University, Miami, FL, USA; Psychological Brain Sciences, Indiana University, Bloomington IN, USA; GIGA Cyclotron Research Centre in vivo imaging, University of Liege, Liege Belgium; Donders Institute for Brain, Cognition and Behaviour, Radboud University, Nijmegen, NL; NatMEG, Karolinska Institutet, Stockholm, Sweden; Department of Radiology and Biomedical Imaging, Yale University, New Haven, USA; Molecular Imaging Branch, National Institute of Mental Health, National Institutes of Health, Bethesda, USA; Department of Computer Science, University of Copenhagen, Copenhagen, Denmark

## Abstract

The Brain Imaging Data Structure (BIDS) is a standard for organizing and describing neuroimaging datasets. It serves not only to facilitate the process of data sharing and aggregation, but also to simplify the application and development of new methods and software for working with neuroimaging data. Here, we present an extension of BIDS to include positron emission tomography (PET) data (PET-BIDS). We describe the PET-BIDS standard in detail and share several open-access datasets curated following PET-BIDS. Additionally, we highlight several tools which are already available for converting, validating and analyzing PET-BIDS datasets.

## Background & Summary

Positron Emission Tomography (PET) was developed in the late 1950s with the ultimate goal of measuring and visualizing physiological processes *in vivo* such as metabolism, blood flow and the concentration of proteins in various receptor systems^1–3^. Since then, PET has been used extensively in pre-clinical and clinical settings, mostly for oncological purposes^4^, but increasingly also in areas of cardiology and neurology (for investigation of, e.g., synaptic plasticity, neuroinflammation, and neurodegeneration^5^). For brain imaging specifically, PET has largely been applied together with high affinity radiolabeled molecules to quantify the brain’s distribution, concentration and drug occupancy of proteins, using pharmacokinetic models^6^. The outcomes of this work have provided significant insights into the complex neurobiology of receptor systems in the healthy and diseased brain, as well as advancements in our understanding of pharmacological treatments^6^. However, compared to imaging modalities such as structural Magnetic Resonance Imaging (MRI), experiments using PET often involve data from multiple sources including, e.g., chemical characteristics of radiolabelled molecules and blood and metabolite data acquired during the imaging procedure (Figure 1). The acquisition and availability of different types of data related to the PET experiment depends on the biological target of interest, and complicates the standardization of the acquired data for storage, analysis and sharing. Furthermore, due to differences in PET scanner vendors and blood sampling devices across PET centers, data are often stored in different file formats, and with varying nomenclature describing the same type of data. Historically, the field of PET brain imaging attained maturity with the adoption of a consensus nomenclature for pharmacokinetic modeling of PET data in 2007^7^. Recently in 2020, similar to the Organization for Human Brain Mapping recommendations from the Committee on Best Practices in Data Analysis and Sharing (COBIDAS) for MRI and MEEG, consensus guidelines for describing the content and format of PET brain data in publications and archives has emerged^8^.

**Figure 1.**
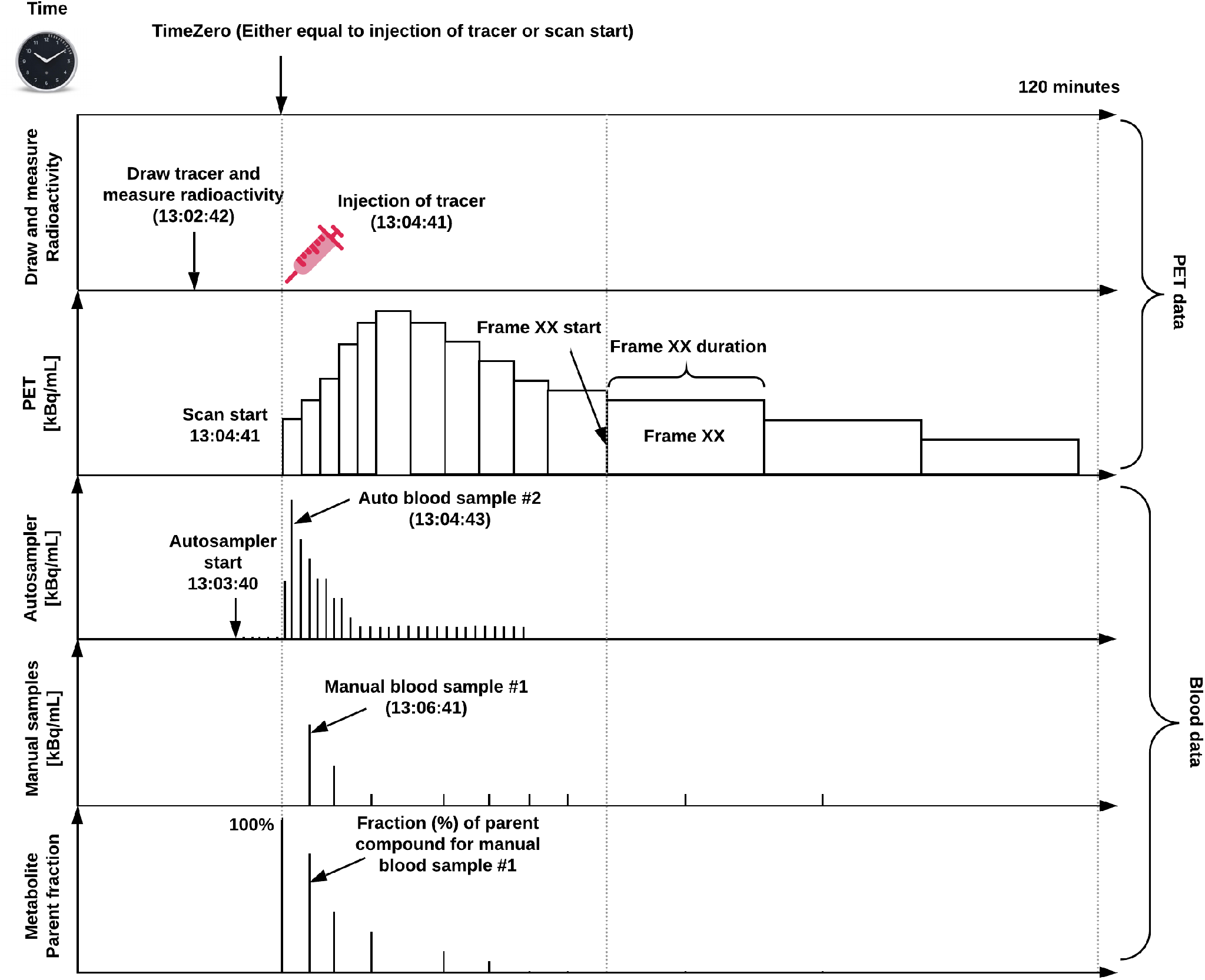
Overview of a common PET experiment, including blood measurements, and defined on a common time scale. Note, “time zero” can either be defined as time of injection or scan start, and all the PET and blood data should be decay-corrected to this time point.

The Brain Imaging Data Structure (BIDS) standard was originally developed for MRI as a community standard for organizing and sharing brain imaging study data within and between laboratories^9^. While the focus was originally targeting MRI, and especially structural and functional MRI (fMRI), BIDS has since rapidly expanded to include many different imaging modalities, including magnetoencephalography (MEG)^10^, electroencephalography (EEG)^11^, intracranial electroencephalography (iEEG)^12^, and linking brain imaging to genetics^13^. BIDS mainly addresses the heterogeneity of data organization by following the FAIR principles (findability, accessibility, interoperability, and reusability)^14^. Findability and reusability are addressed in BIDS by providing rich metadata in dedicated sidecar files. Interoperability is addressed by using the existing Neuroimaging Informatics Technology Initiative (NIfTI) format for storing brain imaging data, text files arranged according to the JavaScript Object Notation (JSON) format for complementary metadata, and Tab Separated Value (TSV) format for tabular data. While accessibility is not directly addressed within the BIDS standard itself, the existence of such a standard facilitates the development of public data repositories. The largest of these repositories, OpenNeuro (https://openneuro.org), is already built around the BIDS standard, and a new repository fully dedicated to PET scans (OpenNeuroPET) is currently under development (https://openneuro.org/pet). BIDS also fosters interoperability and reuse of already acquired data by defining how to structure and store data using naming conventions and dedicated metadata files. Because BIDS data follow a common data structure and description, the proliferation of BIDS datasets incentivizes the creation of analysis pipelines that target this structure, the adoption of which promotes verifiable and reproducible research practices.

In this work, we present the main features of the extension of BIDS to include PET data (PET-BIDS). This extension largely builds upon the original BIDS specification^9^, and the guidelines for the content and format of brain PET data in publications and archives^8^. The full documentation of the PET-BIDS extension can be found in the general BIDS specification ^1^.

## PET-BIDS summary

The extension of BIDS to include PET data largely aligns with the BIDS specification for other imaging modalities, describing a way for organizing PET data and specifying metadata for PET experiments. A rough overview of the directory structure is given in Figure 2, and a detailed example is given in Figure 3. Each subject’s data corresponds to a directory which contains subdirectories for each acquisition session (e.g., *baseline*, *rescan* or *intervention*) and imaging modality (e.g., *pet*). The subdirectories are also accompanied by a `dataset_description.json` file containing generic information about the dataset, providing full credit to the authors sharing the data. Within each subject directory, the `/pet` subdirectory contains the PET imaging data and the corresponding metadata. PET imaging data are stored in 4D (or 3D if only one volume was acquired) NIfTI files with a `_pet` suffix. When acquiring several volumes (frames in PET terminology) these should be stored in 4D in chronological order (the order they were acquired in). The imaging data are accompanied by a `_pet.json` sidecar file which details the metadata of the PET acquisition. All the metadata are in accordance with the guidelines for the content and format of brain PET data^8^.

**Figure 2.**
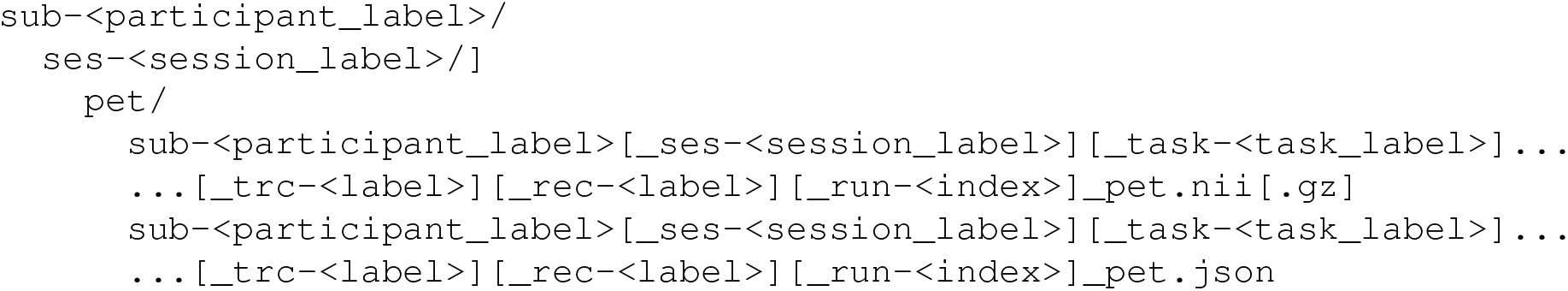
The template for the naming convention of the PET-BIDS file structure. The naming convention follows a key-value pair defining various steps of the acquisition, including the session, task, acquisition, reconstruction, and run. The main imaging data file has a `_pet` suffix stored in the NIfTI format (*.nii). The imaging data are accompanied by a JSON sidecar file containing all the necessary metadata needed to understand the PET data.

**Figure 3.**
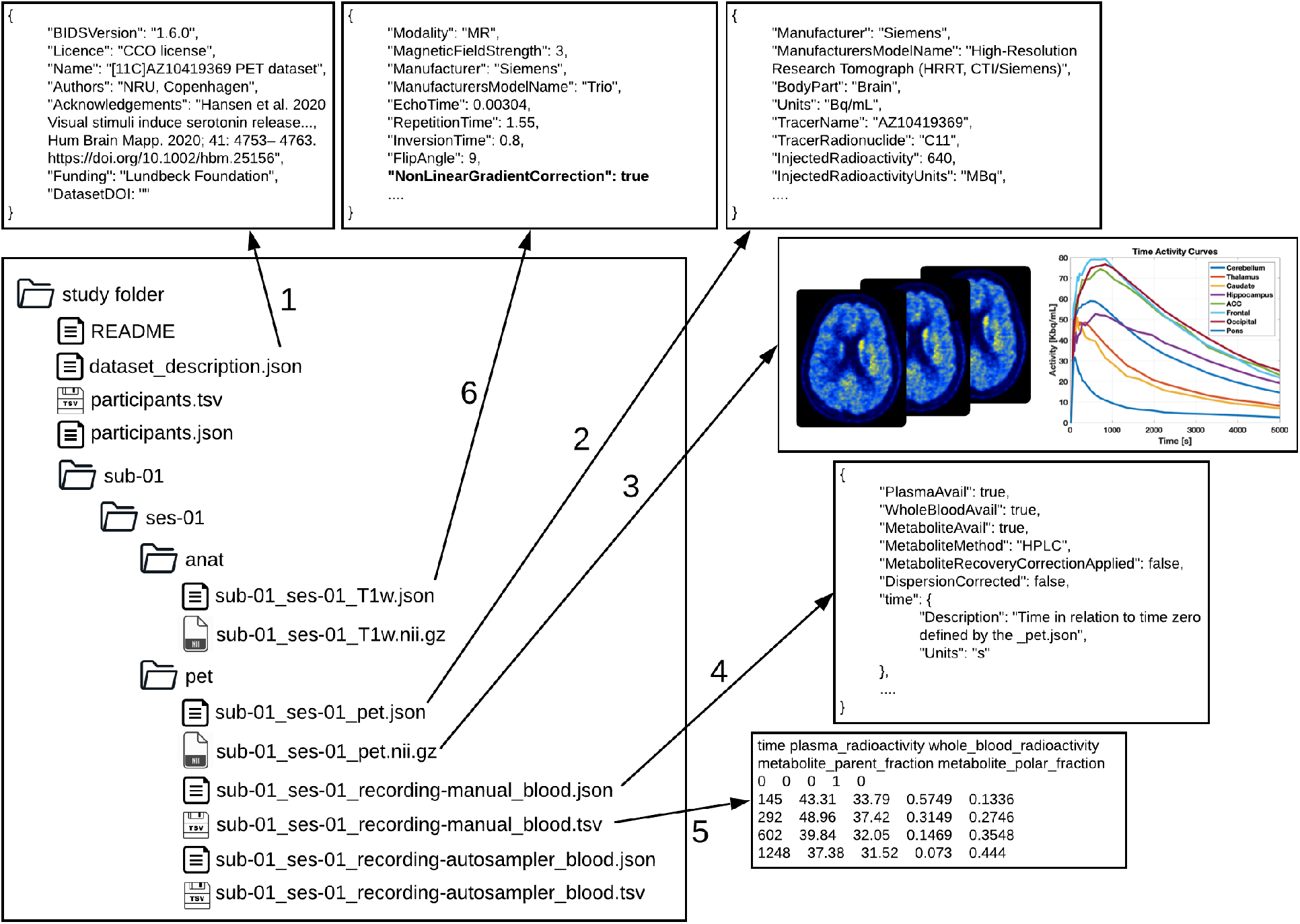
Exemplary PET-BIDS dataset with a dataset description, including adequate acknowledgements (1), previews of PET files (2,3), including blood (4,5) and MRI data (6). The left side shows a directory tree of a common PET-BIDS dataset, with files in the root directory describing the dataset (`README` and `data_description.json`), a file with participant-specific information (`participants.tsv`), and a JSON sidecar file describing the metadata needed to understand the corresponding TSV file. Next to the files in the root directory, there are subject directories named `sub-<label>` for each study participant. In the subject directory lies all acquired data divided into modalities (anat and pet, for the PET and structural MRI, respectively). The content of the pet directory are displayed in the right side of the figure, including the metadata of the raw PET data (2), and the associated imaging data (3). The metadata for the blood data acquired using manual sampling is stored in a JSON sidecar file (4), with the corresponding blood specified in the TSV file (5). The columns in the TSV file contain 1) time, 2) plasma radioactivity, 3) whole blood radioactivity, 4) metabolite parent fraction, and 5) metabolite polar fraction. Blood data acquired using an autosampler is also available following a similar structure as (3) and (4). The PET data may be accompanied with MRI data for co-registration and region definition (6). In this case, it is required to specify if the MRI data has been corrected for gradient non-linearities (*NonLinearGradientCorrection*) to allow for correct co-registration with the PET data.

The file naming structure for PET data closely follows the general BIDS guidelines^9^, using specified key-value pairs joined by hyphens and separated by underscores. Multiple sessions (typically visits) are encoded by adding an extra layer of directories in the form of `ses-<label>`. Hence, a single session study, `sub-<label>` would have a subdirectory `/pet` which contains PET files using the naming pattern `sub-<label>_ses-<label>_pet.nii[.gz]` corresponding to several acquisitions of PET data. The session label should be used in cases such as test-retest or baseline-intervention setups. Additionally, a `task-<label>` can be inserted in a similar way as task-based and resting state Blood Oxygen Level Dependent (BOLD) fMRI data in the existing BIDS standard. For example, in the case of studies using combined PET/fMRI, subject-specific tasks may be carried out during the acquisition within the same session. Therefore, it is possible to specify `task-<label>` in accordance with the fMRI data. Specifically for PET, multiple acquisitions per subject using different tracers during the same session are possible and the `trc-<label>` must be used to distinguish between different tracers. Please keep in mind that the label used is arbitrary and each file requires a separate JSON sidecar file with details of the tracer used. Also, specifically for PET a reconstruction key `rec-<label>` can be used to distinguish different types of reconstructions of the PET data. The `rec-<label>` has four reserved values: *acdyn*, for reconstructions with attenuation correction of dynamic data; *acstat*, for reconstructions with attenuation correction of static data; *nacdyn*, for reconstructions without attenuation correction of dynamic data; *nacstat*, for reconstructions without attenuation correction of static data. Further information about the reconstruction should be added to the accompanying `_pet.json` metadata file. Finally, the `run-<index>` can be used if one scan type/contrast is repeated multiple times within the same scan session/visit. For example, for dynamic PET acquisitions, subjects may have to leave the scanner to use the bathroom. While leaving the scanner would interrupt an MR acquisition, in PET this disruption is more appropriately considered missing data during a run, and the acquisition would still be considered the same session/run. However, there are also cases of acquisitions where this definition might not be entirely clear, and it will be up to the researcher to decide what makes most sense. For example, dual-time-window acquisitions^15^ could be considered two runs within the same session, but it could also be considered a single run with missing data between the two time windows.

### Blood data availability

If blood data are available, such as arterial or venous samples acquired during the PET experiment, they are stored in the `/pet` folder alongside the corresponding PET data (Figure 3). Blood can be sampled by an autosampler, for continuous monitoring of whole blood radioactivity, and/or manually drawn for discrete blood samples. Therefore, the recording key `recording-<label>` for blood data has two reserved values: 1) *autosampler*, and 2) *manual*. The blood metadata should be stored in a JSON sidecar file with a `recording-<label>` and a `_blood` suffix, containing information about what blood data are available (e.g. radioactivity in plasma and/or whole blood and parent compound). The blood JSON sidecar file should be accompanied by a tabular TSV file with similar naming convention, containing all the values of the available blood data. All blood data should be reported according to a unique reference time-scale in relation to a predefined time zero defined by the PET data (Figure 1). The definition of time zero will be further explained below.

## Specific PET-BIDS considerations

In order to construct the `_pet.json` sidecar file which details the PET experiment metadata, a description of a common PET experiment is necessary. In Figure 1 we present an overview of a common PET experiment (which includes the sampling of blood data, including plasma, whole blood and metabolite data). The experiment is defined on a single time scale relative to a predefined “time zero”. Notably, “time zero” will often be defined as time of injection or scan start, and the injected dose, the PET data, and blood data should optimally all be decay-corrected to time zero. However, because the time of injection does not always coincide with scan start, PET data may not be decay-corrected to the time of injection. Whether the image has been decay-corrected may be indicated in its metadata (using the fields: *ImageDecayCorrected* and *ImageDecayCorrectionTime*). The flexibility in choice of time zero to either scan start or injection time was chosen to maximize ease of use and adoption of PET-BIDS by the broadest possible spectrum of the PET community, due to potentially large differences in experimental design between PET studies. For example, scan start and injection time may not always coincide, and due to radioactive decay of the radiotracer, it is important to be aware of post-hoc decay correction. Importantly, the injected dose should always be decay corrected to the time of injection.

Across the diverse set of radiotracers and experimental designs in PET, it will not always be possible to enter the required metadata following the guidelines for sharing of PET data^8^. For example, while the injected mass and specific radioactivity are required metadata according to the guidelines^8^, this is not possible to measure for certain radiotracers such as [^18^F]FDG due to its mass being too low to measure. In these cases, the values for injected mass and specific activity may be set to “n/a” to indicate missing values. We note that for required metadata, this is currently only valid for injected mass and specific activity, although future releases of the PET-BIDS specification may tackle further challenges in use cases that deviate from the current guidelines.

In the case of including MRI data with PET data, it is necessary to pay specific attention to the format the MR images are in, such as whether they have been unwarped to correct for gradient non-linearities. There is a specific metadata field in the BIDS specification for MRI^9^ named *NonlinearGradientCorrection* which indicates this (please see Figure 3 for an example). The main reason for the importance of this is that the MRI needs to be corrected for nonlinear gradients causing spatial distortions in order to have the same shape as the accompanying PET scans for co-registration^8^. Therefore, it is required to specify whether the corresponding MR images have been corrected for gradient non-linearities, using the *NonLinearGradientCorrection* metadata field, if PET data are present.

In general, SI units must be used to describe the data such as “Bq/mL” for radioligand concentration, and seconds for time measurements relative to either scan start or injection time (“time zero”).

## Public PET-BIDS datasets

Several example datasets (with zero-byte, i.e., empty, NIfTI files) are publicly available in the BIDS-examples GitHub repository (https://github.com/bids-standard/bids-examples). The first two of these datasets (full version) are also openly available on OpenNeuro formatted using the PET-BIDS standard:

- The CIMBI Database [^11^C]DASB PET Example Dataset consists of test and retest measurements from two individuals to measure serotonin transporter availability^16^. No blood data are available for this dataset. It was collected as a part of the CIMBI database (https://doi.org/10.18112/openneuro.ds001420.v1.0.1)^17^.
- The NRM2018 Grand Challenge dataset consists of baseline and intervention data from five individuals. No blood data are available for this dataset (https://doi.org/10.18112/openneuro.ds001705.v1.0.1)^18^.
- The CIMBI Database [^11^C]CIMBI-36 PET Example Dataset consists of a single dynamic PET measurement of a pig to measure serotonin 2A receptor availability. Blood and metabolite data are available for this dataset. It was collected as a part of the CIMBI database (https://github.com/bids-standard/bids-examples/tree/master/pet001).
- The CIMBI Database [^11^C]CIMBI-36 Intervention Example Dataset consists of a single dynamic PET measurement of a pig using bolus-infusion, and with a pharmacological ketanserin intervention during the scan. Blood and metabolite data are available for this dataset (https://github.com/bids-standard/bids-examples/tree/master/pet004).
- The CIMBI Database [^11^C]AZ10419369 Visual Stimuli Example Dataset consists of two dynamic PET measurements of a single subject using combined PET/MRI. The first scan is a baseline scan, whereas the second scan includes a visual stimuli task during the scan. No blood data are available for this dataset. It was included in Hansen et al. 2020^19^ (https://github.com/bids-standard/bids-examples/tree/master/pet005).

## Community tools for data sharing and analysis

### The BIDS validator

Data curated into the PET-BIDS standard can be validated for BIDS compliance by using the “bids-validator”^20^, a JavaScript application checking for the completeness and consistency of the data. The BIDS validator runs locally as a command line version (via Node.js), as a Docker container, or as a browser-based application (https://bids-standard.github.io/bids-validator/). Using this important validation software, PET researchers are provided with feedback about incompatibility errors as well as warnings if important pieces of metadata are missing. In addition to providing a version of the validator as open source software, we are collaborating with the software developers of major PET analysis tools (PMOD, SPM, MIAKAT and PETsurfer^21^) to facilitate rapid adoption and support of this format, and working with major PET centres to help PET researchers convert their data into PET-BIDS format. Several software tools already exist to convert dicom files and ECAT data into BIDS format, such as dcm2niix^22^ (https://github.com/rordenlab/dcm2niix), however, the output may need *post-hoc* editing if required metadata are not available in the imaging header files.

### The BIDS starter kit

The BIDS starter kit is a tool to help researchers get started with the BIDS data structure (https://github.com/bids-standard/bids-starter-kit). It consists of a collection of community-driven guides, tutorials, helper scripts, and wiki resources. A tutorial that describes how to create a BIDS-compatible PET data set has been provided on the starter-kit wiki (https://github.com/bids-standard/bids-starter-kit/wiki), and MATLAB (bids-matlab; https://github.com/bids-standard/bids-matlab) and Python (pybids; https://github.com/bids-standard/pybids) packages are also available to produce and/or work with PET sidecar JSON and TSV files. These packages are freely available on GitHub.

### Sharing of acquired PET data

According to the guidelines for the content and format of brain PET data^8^, acquired PET data are defined as PET data after reconstruction into 3D or 4D frames. These data may be shared in repositories such as OpenNeuro (https://openneuro.org), which is an open archive for analysis and sharing of public neuroimaging data spearheading the movement of best practices within MRI, MEG, EEG, iEEG, ECoG, ASL and now PET^23^. The OpenNeuro platform is the successor of OpenfMRI (established in 2011, https://openfmri.org/), and the project enjoys relatively wide acceptance by the field and capitalizes on the BIDS standard. It has been running for almost a decade and is one of the fastest growing image databases, with about 12 new datasets being added per month. All datasets on OpenNeuro are validated for BIDS compliance prior to upload.

### Data analysis pipelines and sharing of derived PET data

BIDS also offers the possibility to build fully reproducible analysis workflows using the concept of BIDS applications^24^. A BIDS application is a software container capturing all the dependencies of a neuroimaging analysis pipeline (e.g., fMRIprep) that takes a BIDS formatted data set as input. Each BIDS application has the same core set of command line arguments, making them easy to run and integrate into automated platforms, allowing for full computational reproducibility. Several open-source initiatives for PET are currently under development to BIDS applications, including PETSurfer (https://surfer.nmr.mgh.harvard.edu/fswiki/PetSurfer), APPIAN (https://github.com/APPIAN-PET/APPIAN) and kinfitr (https://github.com/mathesong/kinfitr) providing tools to carry out preprocessing and/or pharmacokinetic modeling of PET data. We also highly recommend the great resources by the TURKU PET centre (http://www.turkupetcentre.net/), which have been providing thorough documentation and analysis tools for PET for several decades. The resulting outputs from BIDS applications can be shared using BIDS derivatives standards, describing the outputs of common preprocessing pipelines and pharmacokinetic models. The specification for PET-BIDS derivatives (BIDS Extension Proposal 23) is currently under development, and will capture data and metadata sufficient for a researcher to understand and reuse the output of a common PET analysis pipeline, including preprocessing and pharmacokinetic modeling.

## Conclusion

The PET extension to BIDS specifies a structured way of storing PET data and metadata. BIDS is a community-driven project, and the PET-BIDS specification was created in a joint effort made by the PET community (open and with community peer review) aligning with the consensus guidelines for the content and format of PET brain data in publications and archives^8^. PET-BIDS will make data sharing and software development easier, facilitate rapid development, adoption and application of new tools and methods, and ultimately foster collaboration between researchers to combine data sets from different centers to achieve larger sample sizes and improved statistical power to test hypotheses.

## Acknowledgements

We are grateful to the Neurobiology Research Unit, Copenhagen, Denmark, and the Centre for Neuroimaging Sciences, IoPPN, Kings College London, London, UK, for providing the data presented in this manuscript, including uploading it to OpenNeuro. This work was supported by the Novo Nordisk Foundation (NNF20OC0063277). MN was supported by the Independent Research Fund Denmark (DFF-0129-00004B), and MG was supported by the Elsass foundation (18-3-0147). AG is supported by the KCL funded CDT in Data-Driven Health and this represents independent research part funded by the National Institute for Health Research (NIHR) Maudsley Biomedical Research Centre at South London and Maudsley NHS Foundation Trust and King’s College London and part funded by GlaxoSmithKline (GSK). FF, CM, and RB were supported by the National Institute Of Mental Health of the National Institutes of Health (R24MH117179).

## Author contributions statement

MG and MN: Conception and design of the specification, moderating community interactions, coding of the bids-validator extension, preparation of datasets and examples, writing the manuscript. GJM: critical review and editing of the specification, coding of PET-BIDS tools (KinFitr), final review of the version submitted. HDH, AT, GS, GR, MV, AG, MY, MT, TF, AG, HB, AR, JRD, TB, FF, CJM, KJG, RWB, SA, RG, TS, GN, CP, CP, RO, JG, REC, GMK, RBI: critically reviewed the specification and final approval of the version submitted.

## Footnotes

1 https://bids-specification.readthedocs.io/en/stable/

